# The PP1 phosphatase exhibits pleiotropic roles controlling both the tachyzoite cell cycle and amylopectin-steady state levels in *Toxoplasma gondii*

**DOI:** 10.1101/2023.08.10.552784

**Authors:** Asma Sarah Khelifa, Tom Boissavy, Thomas Mouveaux, Tatiana Araujo Silva, Cerina Chhuon, Marcia Attias, Ida Chiara Guerrera, Wanderley De Souza, David Dauvillee, Emmanuel Roger, Mathieu Gissot

## Abstract

Virulence of apicomplexan parasites is based on their ability to divide rapidly to produce significant biomass. The regulation of their cell-cycle is therefore key to their pathogenesis. Phosphorylation is a crucial post-transcriptional modification that regulates many aspects of the eucaryotic cell cycle. The phosphatase PP1 is known to play a major role in the phosphorylation balance in eukaryotes. We explored the role of TgPP1 during the cell cycle of the tachyzoite form of the apicomplexan parasite *Toxoplasma gondii*. Using a conditional mutant strain, we show that TgPP1 regulates many aspects of the cell cycle including the proper assembly of the daughter cells inner membrane complex (IMC), the segregation of organelles and nuclear division. Unexpectedly, depletion of TgPP1 also results in the accumulation of amylopectin, a storage polysaccharide that is normally found in the latent bradyzoite form of the parasite. Using transcriptomics and phosphoproteomics, we show that TgPP1 mainly acts through post-transcriptional mechanisms by dephosphorylating target proteins including IMC proteins. TgPP1 also dephosphorylate a protein bearing a starch binding domain. Mutagenesis analysis reveals that the targeted phospho-sites are linked to the ability of the parasite to regulate amylopectin steady-state levels. Therefore, we show that TgPP1 has pleiotropic roles during the tachyzoite cell cycle regulation, but also regulates amylopectin accumulation.

## INTRODUCTION

Apicomplexa is a phylum that comprises single-celled, obligate, intracellular protozoan parasites. Within this phylum, there are several species of human pathogens, such as *Plasmodium* spp. (the causative agent of malaria), *Toxoplasma* (which causes toxoplasmosis), and *Cryptosporidium* spp. (the cause of cryptosporidiosis). Interestingly, *T. gondii* has emerged as a model for other apicomplexan parasites due to its genetic tractability. Acute toxoplasmosis is caused in intermediate host by the rapid proliferation of the tachyzoite form. Chronic toxoplasmosis is due to the establishment of the encysted form of the parasite (bradyzoites) in specific tissues (mainly brain and muscles). The establishment of the bradyzoite cysts in neurons or muscle cells is of interest in terms of pathology in humans, since reactivation of the cysts can lead to the deadly form of the disease (cerebral toxoplasmosis). It is also crucial for transmission since cysts contained in muscles are one of the main infective routes for humans through the consummation of undercooked meat. Therefore, the ability to rapidly proliferate as a tachyzoite and to switch to the latent bradyzoite form are key elements of pathogenesis in humans and other intermediate hosts.

Diseases triggered by apicomplexan parasites involve an uncontrolled increase in parasite numbers, leading to inflammation and destruction of host cells. Although these parasites undergo a sexual cycle in the definitive host, the pathogenesis is primarily driven by asexual replication cycles occurring within the host’s cells. Asexual cycles involve a tightly regulated control of the cell cycle, ultimately resulting in the production of new daughter cells containing one nucleus and a complete set of organelles ^1^. Although regulation of the cell-cycle involves transcriptional control of gene expression ^2^, post-transcriptional regulation of critical mechanisms has been shown to occur in these parasites ^1^. Among these post-translational modifications phosphorylation and dephosphorylation regulate important molecular functions including parasite cell division ^3,4^ and both *T. gondii* and *P. falciparum* global phosphoproteome show extensive phosphorylation of a large proportion of proteins ^5^ suggesting an important contribution of these post-transcriptional marks in the life cycle of these parasites.

*T. gondii* kinases have been demonstrated to have crucial roles in the division of the tachyzoite. The proper division of the centrosome is regulated by TgCDPK7 ^6^, TgMAPK-L1 ^7^ and TgMAPK2^8^. Moreover, cell division of the *T. gondii* tachyzoite is regulated by five cyclically expressed kinases denoted as TgCrks; these kinases control several cell cycle checkpoints and ensure the smooth progression of the cell cycle ^9^. Only a handful of protein phosphatases have been characterized in depth in this parasite and many protein phosphatase functions remain to be unraveled ^4^. Seventeen serine/threonine protein phosphatases were mutated using the CRISPR/Cas9 system and only TgPP7 demonstrated a prominent role in virulence ^10^. TgPP2A has been shown to play a role in differentiation from tachyzoite to the bradyzoite stage by regulating starch metabolism ^11,12^, a process that is also controlled by the kinase TgCDPK2 ^13^. Moreover, TgPP1, a serine/threonine phosphatase that has been extensively studied in higher eukaryotes, has been shown to play a critical role in promoting motility of the parasite in response to Ca^2+^ upon egress from the host-cell ^14^. Notably, upon the release of Ca2+ from intracellular stores, TgPP1 exhibited relocalization to the apex of the parasite ^14^. Phosphoproteomics analysis after treating parasites with zaprinast (a cGMP activator that ultimately stimulates egress of the parasite), revealed that TgPP1 is likely involved in dephosphorylating crucial targets during egress and motility ^14^. The affected pathways included transmembrane transport, cyclic nucleotide synthesis, ubiquitin transfer, and cytoskeletal motor activity ^14^, revealing the crucial roles of TgPP1 in the control of the ultimate steps of the intracellular cycle of the parasite. However, the role of TgPP1 during the cell cycle and in controlling intracellular growth has not been explored. Indeed, TgPP1 forms a complex with TgLRR1^15^ and TgI2^16^, two homologs of known regulators of cell cycle function in other eukaryotes ^17^, suggesting a role in controlling the cell cycle in this parasite.

In this study, we produced an inducible knock-down mutant of TgPP1 and characterized its phenotypes during the tachyzoite cell-cycle. We demonstrate that TgPP1 has a crucial role in regulating the multiple pathways that are essential for the production of daughter cells such as organelle division and segregation. TgPP1 also ensures normal assembly of the daughter cell inner membrane complex. Phospho-proteomics demonstrated the differential phosphorylation of several IMC proteins and other targets that are linked to the regulation of the cell cycle. Moreover, we demonstrated the role of TgPP1 in starch production and identified an unknown protein that is crucial for the balance of starch production and whose activity is, at least partially, regulated through dephosphorylation by TgPP1.

## MATERIALS AND METHODS

### Parasite culture, transfection, and purification

*Toxoplasma gondii* tachyzoites belonging to the Type I RH Δ*ku80* Tir1 strain were grown *in vitro* in human foreskin fibroblast (HFF) cells using Dulbecco’s Modified Eagles medium supplemented with 10% fetal bovine serum (FCS), 2mM glutamine, and 1% penicillin-streptomycin. The Type I Δ*ku80* Tir1strain has the *Ku80* gene deleted with the aim of promoting homologous recombination (Huynh & Carruthers, 2009). In addition to expressing the Transport Inhibitor Response (Tir1) protein which has a role in the inducible degradation of targeted proteins using an Auxin-inducible degradation (AID) system (Brown et al., 2018). Parasite strains were grown in T25 ventilated flasks containing HFF cell monolayers and maintained within a HERAcell VLOS 160i CO_2_ incubator (Thermo Scientific) at 37 ^◦^C and 5% CO_2_. Electroporation was used to introduce transgenes into the parasite’s genome by using a BTX Electro Cell Manipulator 600 at 1.5 kV.cm^-1^, 25 µF capacitance, and 24 Ω resistance. Transgenic parasites were selected by using 25 ug/ml mycophenolic acid (MPA) and 50ug/ml xanthine. Parasite clones were produced by using limiting dilution. Experiments consisting of DNA, total RNA, and protein purification involved tachyzoite collection by mechanical lysis using a 17-guage needle followed by a 24-guage needle (Terumo AGANI) followed by the filtration of the lysate using a 3um polycarbonate membrane (Whatman).

### Generation of transgenic T. gondii strains

The iKD TgPP1 line was generated by utilizing the Type I RH Δ*ku80* Tir1 strain as a parental strain. A CRISPR/Cas9 plasmid containing a gRNA targeting the 5’ end of the endogenous *TgPP1* gene (TGGT1_310700) and a PCR product consisting of the HXGPRT selection cassette (HXGPRT-T2A-AID-Ty) flanked by 30 bp of homology were used for transfection. The PCR product contained a Skip peptide (T2A) allowing for the dual expression of protein under a single promoter followed by cleavage of the TgPP1 protein from the HXGPRT protein in order to promote the degradation of the TgPP1 protein by the proteasome once the auxin hormone is added. The PCR product was amplified by using the pHXGPRT-2TA-AID-Ty plasmid as a template. Ten µg of PCR product was transfected with 30µg of pSAG1:: Cas9-U6 targeting the 5’ end of the TgPP1 gene in the RH Δ*ku80* Tir1 strain. Primers used to generate transfection material are included in Supplementary Table 1.

### Generation of trehalose phosphatase mutants

TGGT1_297720 gene was tagged at the 3’end with 3xmyc tag in the RH Δ*ku80* strain using a CrispR/Cas9 expressing construct targeting the 3’ end of the gene. To generate point mutation in the TGGT1_297720 gene, a CrispR/Cas9 construct was design in the 15^th^ intron of the gene. A DNA fragment that included the two phosphosites to be mutated was synthised with a point mutation in the PAM site of the gRNA (located in the 15^th^ intron of the gene). The mutated DNA fragments were by mutagenesis using the Q5 mutagenesis kit to induce point mutations in the codons corresponding to Serine 1054 and Serine 1073. They were mutated to either alanine or aspartate amino acids. These DNA fragments were co-transfected together with the CrispR/Cas9 construct into the RH Δ*ku80* strain. FACS sorting based on the CrispR/Cas9 construct GFP expression was used to isolate the mutant strains and directly clone them into 96 well plates. Each mutant clone was verified using sequencing.

### Growth assays

Growth assays were carried out by the inoculation of 8 x 10^4^ parasites of parental Tir1 and iKD TgPP1 mutant parasites on HFF cell monolayers grown on coverslips in a 24-well plate for 24 hours and 48 hours in the presence and absence of 0.5mM of auxin (AID/indoleacetic acid). Auxin was introduced into the media with the aim of inducing TgPP1 protein degradation. After either 24 hours or 48 hours, infected coverslips were fixated with 4% of paraformaldehyde (PFA). The fixated parasites were stained using anti-TgEno2 in order to stain parasite nuclei as well as anti-TgIMC1 antibodies in order to stain the Inner Membrane Complex (IMC). The number of parasites per vacuole was counted for a total of 100 vacuoles per biological replicate. Each growth assay experiment included a total of three to five biological replicates.

### Plaque assays

In order to carry out the plaque assays, a total of 1000 parasites of either the Parental Tir1 strain or iKD TgPP1 mutant strain were inoculated on a monolayer of HFF cells grown in a 6-well plate either in normal media or media treated with 0.5mM of auxin. Parasites were left to grow for 7 days before fixation using 100% ethanol. Plaques were stained using Crystal Violet for the purpose of plaque visualization. An Excel macro was used to quantify the plaque size for each experimental condition.

### Organelle labelling

Parental Tir1 and iKD TgPP1 parasites were left to grow on monolayers of HFF cells grown on coverslips of 24-well plates in normal media as well as media treated with auxin for 24 hours and 48 hours before fixating parasites with 4% PFA and staining with antibodies. The Inner Membrane Complex (IMC) was labelled using anti-TgIMC1, anti-TgISP1, and anti-TgGAP45. Other organelles consisting of the Golgi, plastid, and centrosome were labelled using anti-TgSORT, anti-TgACP, and anti-TgCentrin1 antibodies, respectively.

### Immunofluorescence assays

Immunofluorescence experiments were carried out after fixation of intracellular parasites grown on coverslips using 4% of PFA for 30 minutes. This was followed by coverslip washing 3 times using 1X PBS buffer. Permeabilization was carried out for 30 minutes using the following buffer (1X PBS, 0.1% Triton 100X, 0.1% glycine, 5% FBS). This was followed by primary antibody incubation for 1 hour. Antibodies were diluted using IFA buffer used for permeabilization. Coverslips containing fixated intracellular parasites were then washed 3 times using 1X PBS. Coverslips were then incubated for 1 hour in DAPI and secondary antibodies coupled to either Alexa-594 or Alexa-488. Coverslips were then washed three times using 1x PBS buffer and mounted onto microscope slides using Moviol. Primary antibodies used included anti-TgIMC1 (a gift from Prof. Ward, University of Vermont), anti-TgEno2, anti-TgISP1 ^18^, anti-TgCentrin1 (a gift from Prof. Gubbels, College of Boston), anti-TgACP (agift from Pr. Striepen, U. Penn), anti-TgSortilin, and anti-TgGAP45 (Prof. Soldati, Geneve University), anti-HA, and anti-Ty (a gift from Dr. Bastin, Institut Pasteur de Paris) were used at the following dilutions: 1:500, 1:1000, 1:500, 1:500, 1:500, 1:500, 1:10000, 1:500, and 1:10000 respectively. Signal was visualized manually by counting 100 parasites for each replicate. A total of three replicates were carried out for each experiment.

Immunofluorescence assay experiments were visualized using the ZEISS LSM88O confocal microscope at a magnification of 63X. Image processing was carried out using the CARL Zeiss Zen software.

### Electron microscopy

iKD TgPP1 parasites were grown on monolayers of HFF cells in normal media or media treated with auxin for 48 hours in T25 flasks. Intracellular parasites were fixated using the following solution: 1% glutaraldehyde in 0.1M sodium cacodylate at a pH of 6.8 and at 4 ^◦^C. Parasite samples were then post-fixated using 1% osmium tetraoxide and 1.5% potassium ferricyanide. This was then followed by using 1% uranyl acetate. Post-fixation was carried out for 1 hour using the following conditions: distilled water, in the dark, and at room temperature. Increasing ethanol concentration solutions were used to dehydrate fixated samples after washing. Epoxy resin was used with the aim of infiltrating parasite samples. This was followed by curation for 24 hrs at 60 °C. Deposition of 70-80 nm-thick sections was carried out in formvar-coated grids. Images were observed using 80kV on a Hitachi H7500 TEM (Federal University of Rio de Janeiro University, Brazil). Acquisition of images was carried out by using a Mpixel digital camera (Federal University of Rio de Janeiro University, Brazil).

### RNA sample preparation and extraction

RNA samples of iKD TgPP1 parasites were collected after inoculating parasite onto monolayers of HFF cells grown in T175 flasks. iKD TgPP1 parasites were grown in normal media (control) as well as media treated with auxin for 24 hrs. RNA extraction was carried out by re-suspending the parasite sample in Trizol (Invitrogen). This was followed by the addition of chloroform (4 ^◦^C) to the sample allowing for the separation of the RNA-containing aqueous phase and the protein-containing organic phase. This was followed by a centrifugation step at room temperature for 10 minutes. The aqueous phase was then transferred into a new tube consisting of cold isopropanol in order to precipitate the RNA followed by a centrifugation step. Washing of the precipitated RNA pellet was carried out using 70% ethanol followed by air-drying the pellet and re-suspension in RNase-free water (Gibco). RNA samples were purified using the RNase-free DNaseI Amplification Grade Kit (Sigma). Extracted RNA quality was verified using an RNA 6000 Nano (Agilent) chip and RNA samples with an integrity score of 8 or greater were used for RNA library preparation.

### RNA-sequencing library preparation

Libraries for the RNA samples were prepared by using the TruSeq Stranded mRNA Sample Preparation Kit (Illumina). Libraries were prepared as per the manufacturer’s instructions. Validation of the libraries was carried out using DNA high sensitivity chips read by an Agilent 2100 Bioanalyzer. Libraries were then quantified by using the KAPA library quantification kit (Illumina) using a 12K QuantStudio qPCR thermocycler.

### RNA-sequencing analysis

RNA libraries were sequenced using a HiSeq 2500 as 50 bp reads by using the sequence by synthesis technique. HiSeq control software and real-time analysis component were used for image analysis. Bcl2fastq 2.17 (Illumina) was used to demultiplex. Dataset quality was verified using FastQC v0.11.8-0. Cutadapt v1.18. was used to treat the adapters for sequencing. Filtering of reads shorter than 50 bp and low-quality bases was carried out by using Trimmomatic v0.38.1. Alignment of datasets using HiSAT2 v2.2.0 was carried out after cleaning of datasets against the *T. gondii* ME49 genome from ToxoDB. Annotated gene expression was quantified using htseq-count from the HTseq suite v0.9.1. DeSeq2 v1.22.1 was used to carry out differential gene expression analysis. P-values were adjusted using the Benjamin-Hochberg method. Adjusted p-values less than 0.05 and a log2 fold change greater than 2 corresponding to differentially expressed genes were kept.

### Phosphoproteomics sample extraction

Mutant iKD TgPP1 parasites were left to grow in T175 flasks before the addition of auxin for 2 hours and 24 hours in T175 flasks. iKD TgPP1 parasites grown in normal media were considered as a control. After parasite purification by filtration and centrifugation, parasite pellet was re-suspended using 8M Urea consisting of Protease and Phosphatase Inhibitor Cocktail (Thermo Fisher Scientific). Parasite was re-suspended in urea solution to a final concentration of 35 million parasites/50 µL of re-suspension solution. Extraction steps were carried out on ice. A Biorupter Sonicator was used at 4 ^◦^C for 10 cycles (30 sec on/off per cycle). This was followed by a centrifugation step for 20 minutes at 4000 rpm at 4 ^◦^C. The supernatant was separated from the pellet and put into a new tube. The Pierce BCA Protein Assay (Life Technologies) was used to quantify the protein concentration of the samples. Samples were then stored at -80 ^◦^C prior to analysis.

### Tryptic digestion

S-Trap^TM^ mini spin column (Protifi, Hutington, USA) digestion was performed on 100 µg of cell lysates, according to manufacturer’s instructions. Briefly, SDS concentration was first adjusted to 5% and samples were reduced with 20 mM TCEP and alkylated with 50 mM chloracetamide for 15 min at room temperature. Aqueous phosphoric acid was then added to a final concentration of 1.2% following by the addition of S-Trap binding buffer (90% aqueous methanol, 100 mM TEAB, pH7.1). Mixtures were loaded on S-Trap columns by 30 sec centrifugation at 4,000 x g. 6 washes were performed before adding the trypsin (Promega) at 1/20 ratio for 2h at 47°C. After elution, peptides were vacuum dried.

### Phosphopeptides enrichment by titanium dioxide (TiO_2_) and phosphopeptides purification by graphite carbon (GC)

Phosphopeptide enrichment was carried out using a Titansphere TiO_2_ Spin tip (3 mg/200 μL, Titansphere PHOS-TiO, GL Sciences Inc, Japan) on 90µg of digested proteins for each biological replicate. Briefly, the TiO_2_ Spin tips were conditioned with 20 µL of solution A (80% acetonitrile, 0,1% TFA), centrifuged at 3,000 x g for 2min and equilibrated with 20µL of solution B (75% acetonitrile, 0,075% TFA, 25% lactic acid) followed by centrifugation at 3,000 x g for 2 min. Peptides were resuspended in 20 µL of 10% acetonitrile, 2% TFA in HPLC-grade water, mixed with 100 µL of solution B and centrifuged at 1,000 x g for 10min. Sample was applied back to the TiO2 Spin tips two more times in order to increase the adsorption of the phosphopeptides to the TiO2. Spin tips were washed with, sequentially, 20 µL of solution B and two times with 20 µL of solution A. Phosphopeptides were eluted by the sequential addition of 50 µL of 5% NH4OH and 50 µL of 5% pyrrolidine. Centrifugation was carried out at 1,000 x g for 5 min.

Phosphopeptides were further purified using GC Spin tips (GL-Tip, Titansphere, GL Sciences Inc, Japan). Briefly, the GC Spin tips were conditioned with 20 µL of solution A, centrifuged at 3,000 x g for 2 min and equilibrated with 20 µL of solution C (0,1% TFA in HPLC-grade water) followed by centrifugation at 3,000 x g for 2 min. Eluted phosphopeptides from the TiO2 Spin tips were added to the GC Spin tips and centrifuged at 1,000 x g for 5 min. GC Spin tips were washed with 20 µL of solution C. Phosphopeptides were eluted with 70 µL of solution A (1,000 x g for 3 min) and vacuum dried.

### nanoLC-MS/MS protein identification and quantification

Samples were resuspended in 42 µL of 0.1% TFA in HPLC-grade water. For each run, 5 µL was injected in a nanoRSLC-Q Exactive PLUS (RSLC Ultimate 3000, Thermo Scientific, MA, USA). Phosphopeptides were loaded onto a µ-precolumn (Acclaim PepMap 100 C18, cartridge, 300 µm i.d.×5 mm, 5 µm, Thermo Scientific, MA, USA) and were separated on a 50 cm reversed-phase liquid chromatographic column (0.075 mm ID, Acclaim PepMap 100, C18, 2 µm, Thermo Scientific, MA, USA). Chromatography solvents were (A) 0.1% formic acid in water, and (B) 80% acetonitrile, 0.08% formic acid. Phosphopeptides were eluted from the column with the following gradient 1% to 40% B (120 minutes), 40% to 80% (1 minutes). At 121 minutes, the gradient stayed at 80% for 5 minutes and, at 126 minutes, it returned to 5% to re-equilibrate the column for 20 minutes before the next injection. Two blanks were run between each replicates to prevent sample carryover. Phosphopeptides eluting from the column were analyzed by data dependent MS/MS, using top-10 acquisition method. Phosphopeptides were fragmented using higher-energy collisional dissociation (HCD). Briefly, the instrument settings were as follows: resolution was set to 70,000 for MS scans and 17,500 for the data dependent MS/MS scans in order to increase speed. The MS AGC target was set to 3.10^6^ counts with maximum injection time set to 200 ms, while MS/MS AGC target was set to 1.10^5^ with maximum injection time set to 120 ms. The MS scan range was from 400 to 2000 m/z. Dynamic exclusion was set to 30 seconds duration.

For total proteomic analysis, peptides were eluted from the column with the following gradient 5% to 40% B (120 min), 40% to 80% (6 min). At 127 min, the gradient returned to 5% to re-equilibrate the column for 20 min before the next injection. MS parameters were the same as those used for LC-MS/MS analysis described above.

### Data Processing Following nanoLC-MS/MS acquisition

The MS files were processed with the MaxQuant software version 1.6.14.0 and searched with Andromeda search engine against the UniProtKB/Swiss-Prot *homo sapiens* database (release February 2021, 20396 entries) and *toxoplasma gondii* strain ATCC 50853 / GT1 (release November 2020, 8450 entries). To search parent mass and fragment ions, we set an initial mass deviation of 4.5 ppm and 0.5 Da respectively. The minimum peptide length was set to 7 amino acids and strict specificity for trypsin cleavage was required, allowing up to two missed cleavage sites. Carbamidomethylation (Cys) was set as fixed modification, whereas oxidation (Met), N-term acetylation and phosphorylation (Ser, Thr, Tyr) were set as variable modifications (only for phosphoproteomics analysis). The match between runs option was enabled with a match time window of 0.7 min and an alignment time window of 20 min. The false discovery rates (FDRs) at the protein and peptide level were set to 1%. Scores were calculated in MaxQuant as described previously ^19^. The reverse and common contaminants hits were removed from MaxQuant output.

The phosphopeptides output table and the corresponding logarithmic intensities were used for phosphopeptide analysis. The phosphopeptide table was expanded to separate individual phosphosites, and we kept all sites identified in all four replicates in at least one group for the 2h condition and in all three replicates in at least one group for the 24h experiment. Missing values were imputed using width=0.3 and down-shift=3. We represented on volcano plots the significantly altered phosphosites (t-test S0=1, FDR=0.05).

For total proteome analysis, we kept only proteins identified in all four replicates in at least one group. Missing values were imputed using width=0.3 and down-shift=2.5. We represented on a volcano plot the significantly altered proteins (t-test S0=1, FDR=0.05).

### Western Blotting

Western blotting was carried out by inoculating 2x 10^6^ parasites of the iKD TgPP1 strain in T175 flasks grown in normal media for 24 hours (control) and grown in media treated with auxin for 1 and 2 hours. Parasite samples were collected by filtration, followed by centrifugation. The obtained pellet was re-suspended in loading buffer consisting of 240 mM Tris-HCl pH 6.8, 8% SDS, 40% saccharose, 0.04% bromophenol blue, and 400 mM DTT. This was followed by a denaturing step by incubating the parasite samples at 95 ^◦^C for 10 minutes. Protein extracts were separated by electrophoresis on an 8% polyacrylamide gel and then transferred onto a nitrocellulose membrane (GE Healthcare) for 90 minutes at 100V. Blocking buffer containing 5% milk in TNT buffer consisting of 100 mM Tris pH 8, 150 mM NaCl and 0.1% Tween was used to block the membrane. Western blot membranes were incubated in primary anti-body for 1 hour, washed 4 times, incubated for another hour in the secondary antibody. Super Signal West Femto Maximum Sensitivity Substrate (Thermo Scientific) was used to reveal protein bands and ChemiDoc^TM^ XRS^+^ (Biorad) was used to visualize protein bands. Antibodies used were anti-Ty used at a dilution of 1/500. Secondary antibody used is species-specific and conjugated to HRP.

### PAS staining

For the PAS staining, Parental Tir 1 and iKD TgPP1 parasites were left to grow on HFF cell monolayers grown on coverslips for 48 hours in the presence and absence of auxin. Periodic acid-Schiff stain was used in order to determine the presence of polysaccharides within the paraites. Visualization of polysaccharides was carried out by using a confocal microscope.

### Amylopectin quantification

Parental Tir1 and iKD TgPP1 parasites were left to grow for 48 hours in the presence and absence of auxin before parasite filtration and centrifugation. Around 200 million parasites were collected for each sample. After purification, parasites samples were stored at -80 ^◦^C and sent for analysis. Purified *T. gondii* cells were resuspended in ice cold phosphate buffered saline (PBS) at a concentration of 20 million parasites per mL. Cell suspensions were then disrupted three times by a French press (13,000 p.s.i.) and centrifuged at 10000g for 30 min at 4°C. The pellets containing amylopectin and cell debris were passed through a self-formed 90% Percoll gradient at 10000g for 30 min at 4°C. The purified amylopectin pellets were washed in ultrapure water, centrifuged twice at 10,000 g and kept dry at 4°C. Polysaccharide amounts were measured by an amyloglucosidase assay using the Enzytec^TM^ starch kit following the manufacturer’s recommendations. In parallel, 30 µg of polysaccharide were resuspended in 20 µl of 100 % dimethyl sulfoxide (DMSO), boiled 10 min and diluted to 10% DMSO. Twenty µl of a freshly prepared iodine solution (0,2% I2; 2% KI) were added to 80 µl of the boiled sample and the absorbance of the complex was monitored from 700 to 400 nm allowing the determination of its λmax (wavelength at the maximal absorbance). Controls including commercial potato amylopectin, potato amylose and rabbit liver glycogen (Sigma) were used and displayed λmax values of 542 nm, 631 nm and 495 nm respectively.

### Statistics

Graph pad Prism software version 8/9 (San Diego, California, USA) was used to analyze all data concerning growth assays, proliferation assays, and ratio counts. Student t-tests were used to determine significant differences between datasets where p values < 0.05 were considered as significant. All experiments were carried out in biological triplicates. For each independent experiment, a total of 100 parasites/vacuoles was counted.

## RESULTS

### TgPP1 is essential for the normal growth and proliferation of *T. gondii* tachyzoites

We generated an inducible knock down mutant of TgPP1(iKD TgPP1) with the aim of determining the role of the TgPP1 protein using the Auxin Inducible Degron (AID) system. The mutant parasite line was produced using CRISPR/Cas9 to insert an *hxgprt-t2a-AID-2ty* insert at the 5’ end of the TgPP1 gene (figure 1a). Previous attempts to generate the transgenic parasite strain by inserting an *hxprt-AID-HA* cassette at the 3’end were unsuccessful. Insertion of the *hxgprt-t2a-AID-2xty* cassette at the appropriate locus within the iKD TgPP1 genome was confirmed by insertion PCR (supplementary figure 1a). The localization of Ty-tagged TgPP1 protein was investigated using IFA and TgPP1 was demonstrated to be localized mainly in the nucleus but also in the cytoplasm when compared with a cytoplasmic marker (TgAlba1^20^) during the tachyzoite intracellular cell cycle (figure 1b). The depletion of TgPP1 was obtained within 1 hour of auxin treatment as verified by Western blot of total protein extracts (figure 1c). A growth assay was carried out for 24 hours in the presence and absence of auxin. We noticed that the growth of the strain was affected in absence of auxin, a phenotype that was also observed for another published TgPP1 conditional mutant ^14^ (supplementary figure 1b). However, the IKD TgPP1 in presence of auxin demonstrated a significant growth defect after 48 hours (figure 1d). We then performed a plaque assay, that measures the ability of the parasite to form plaques in the host cell monolayer over a period of 7 days. In this assay, the iKD TgPP1 was unable to form plaques in the presence of auxin thus demonstrating that the TgPP1 is essential for parasite proliferation as a tachyzoite (figure 1e). This experiment also showed that the size of the plaque produced by the mutant in absence of auxin was smaller than the parental parasite strain, confirming that the insertion of the AID construct was deleterious to the parasite growth (supplementary figure 1c). Overall, these results suggest the essentiality of TgPP1 for the normal growth and proliferation of the tachyzoite.

**Figure 1:**
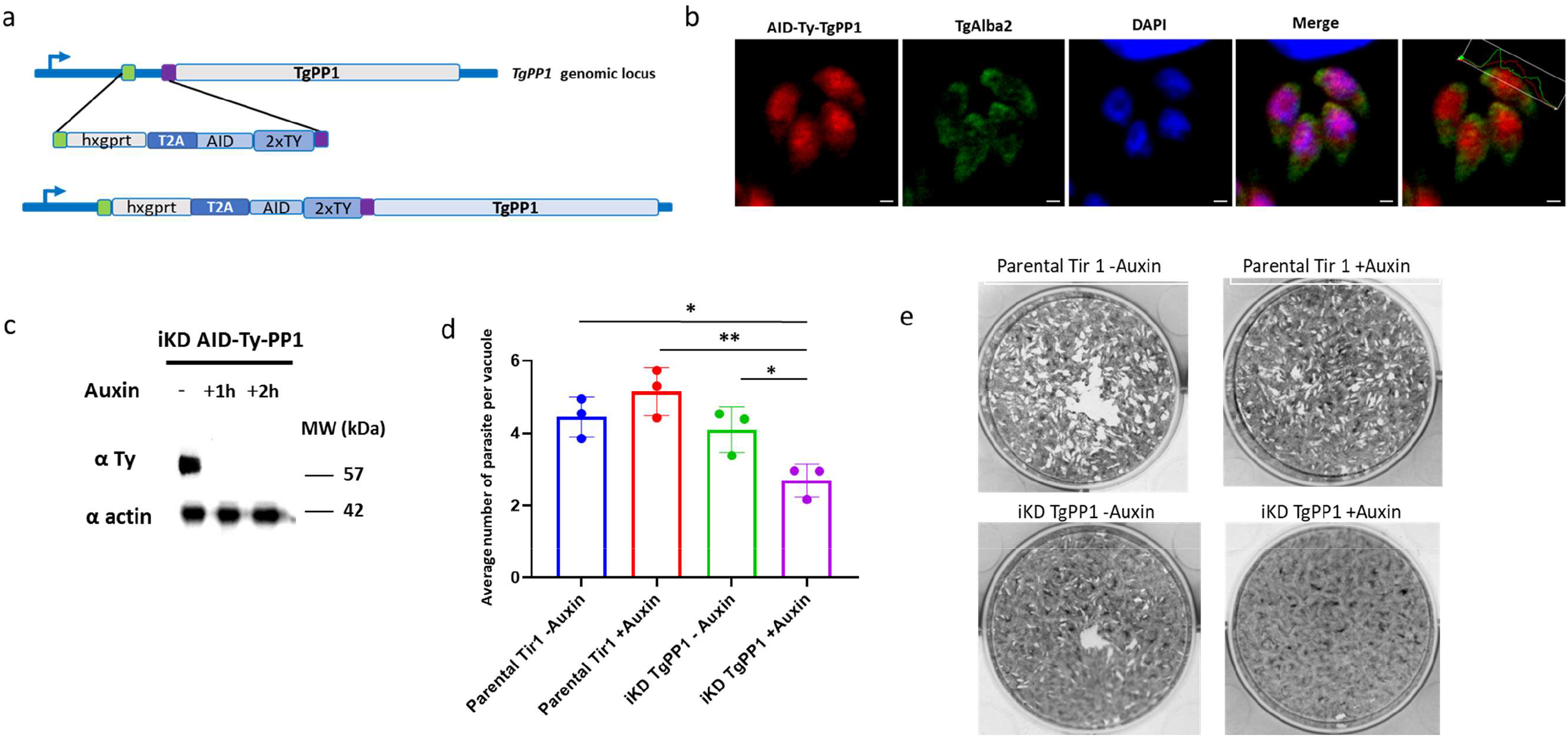
TgPP1 is required for parasite growth and proliferation. (**a)** Schematic representation of the construct used to generate iKD TgPP1 mutant parasites using the Auxin Inducible Degron (AID) system which allows for the inducible degradation of the TgPP1 protein. The system involves introducing the AID domain into the gene of interest. In the presence of Auxin, the AID domain will be recognized by the Tir1 protein and the protein degraded by the proteasome. The HXGPRT-T2A-AID-2Ty cassette is inserted at the 5’end of the gene using the CRISPR/Cas9 gene-editing strategy. **(b)** Confocal microscopy imaging of the AID-Ty iKD TgPP1 parasite in the absence of auxin labelled with anti-Ty (red, TgPP1) and TgAlba2 (green), a protein that localized to the cytoplasm. DAPI (blue) was used to stain the nucleus. The scale bar is indicated in the lower right corner of each image. Measurements of red and green fluorescence throughout length of the parasite is indicated in the upper right corner of the overlayed TgPP1 and TgAlba2 image. **(c)** Western blot of total protein extract from the iKD TgPP1strain in absence and presence of auxin (1 hour and 2 hours) displaying depletion of the TgPP1 protein after the addition of Auxin. Western blots were probed with anti-Ty antibodies to determine the presence of the TgPP1 protein (upper panel). Western blot was normalized using anti-TgActin antibodies (lower panel). **(d)** Growth assay of the Parental Tir1 and iKD TgPP1 strains in the absence and presence of Auxin for 48 hours. The average number of parasites per vacuole was recorded. A Student’s *t*-test was performed. *p< 0.05, **p<0.01; mean ± s.d. (n=3). **(e)** Plaque assay demonstrating proliferation of the Parental Tir 1 and iKD TgPP1 strain in the presence and absence of auxin.

### TgPP1 is important for the formation of the Inner Membrane Complex (IMC)

To better understand the biological role of TgPP1 during the intracellular tachyzoite cell-cycle, we investigated its ability to form the IMC, a double membrane structure located below the tachyzoite’s plasma membrane (PM). Upon depletion of TgPP1, the parasites exhibited an abnormal IMC structure as labeled by an anti-IMC1 antibody, featuring an unstructured network of intertwined IMC1 protein. Upon auxin treatment, the IMC was unable to form properly (figure 2a, lowest panel) as opposed to the IMC structures of the iKD TgPP1 mutant in the absence of auxin treatment as well as the IMC structures of the Parental Tir1 strain (figure 2a, upper and medium panels). The quantification of the percentage of vacuoles with this IMC phenotype demonstrated that 40% of the vacuoles showed a similar phenotype after 48 hours of auxin treatment (figure 2b). This phenotype was accompanied by a missegregation of nuclear material as illustrated in figure 2a (lower panel, iKD TgPP1 + Auxin). Quantification revealed that a similar percentage (40%) of vacuoles showed an important missegregation of nuclear material in the iKD TgPP1 mutant in the presence of auxin (figure 2c) while this phenotype was almost lacking in the parental strain and in absence of auxin. The effect of TgPP1 depletion on additional components of the IMC was also investigated. Upon auxin treatment, TgGAP45, a protein targeted to the plasma membrane and connected with the IMC at the C-terminal end, revealed similar defects in IMC formation in the iKD TgPP1 mutant parasites treated with auxin (supplementary figure 2a). Similar results were obtained with the IMC Sub-compartment protein 1 (TgISP1) which in normal instances localizes to the apical cap of the IMC, and was instead dispersed along the periphery of the iKD TgPP1 parasite in the presence of auxin (supplementary figure 2b). To further confirm the effect of TgPP1 depletion on the IMC structure, electron microscopy (EM) was carried out. In absence of auxin, the iKD TgPP1 mutant parasites showed both the PM and the IMC normally formed and these two structures appeared intact (figure 2d). In presence of auxin, the iKD TgPP1 mutant parasites exhibited an intact and continuous PM along the parasite. In contrast, the IMC was only partially present and discontinuous along the formed PM (black arrows, figure 2e). These EM observations further validate the role of TgPP1 in correctly forming the IMC.

**Figure 2:**
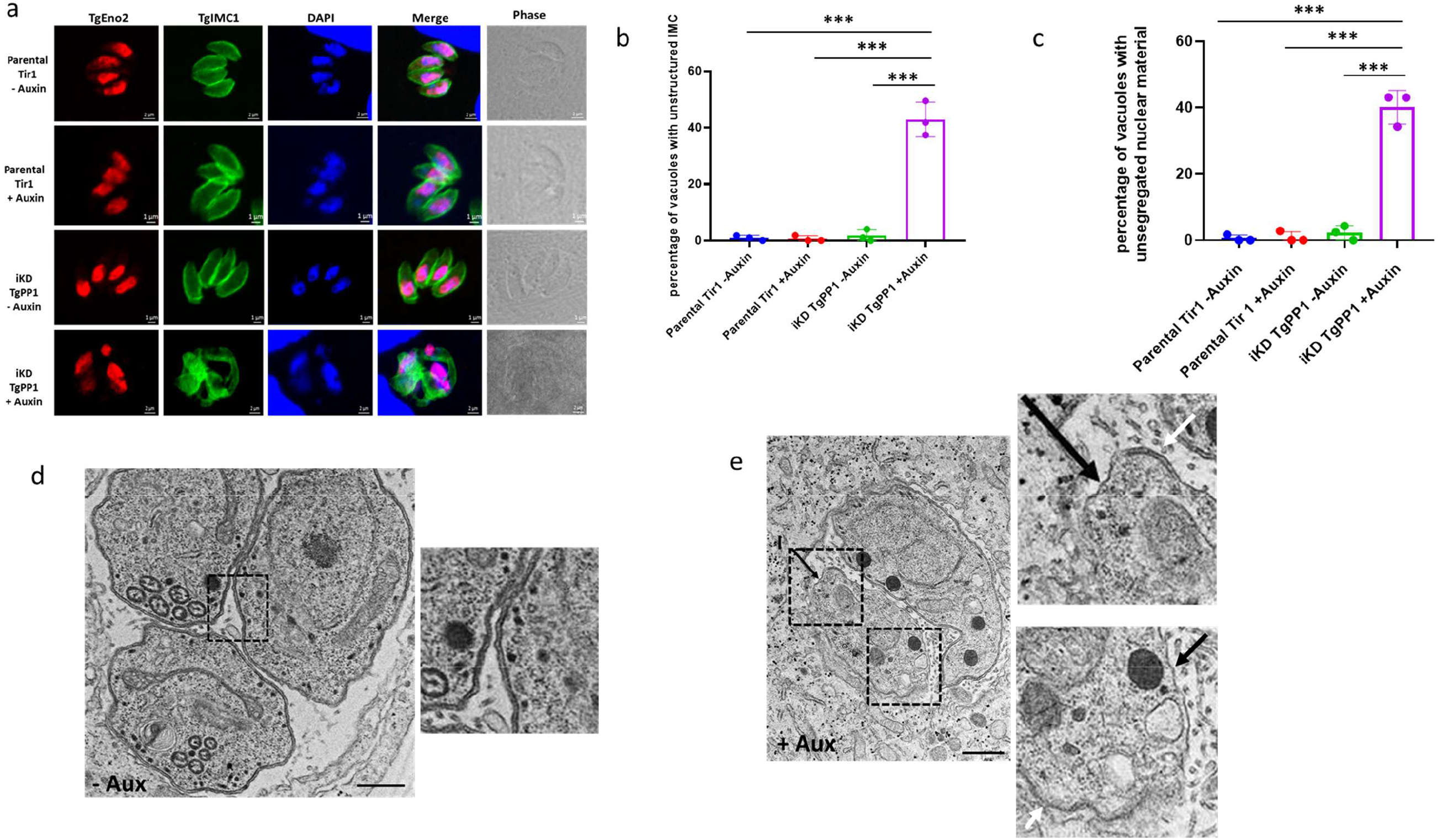
Depletion of TgPP1 results in a collapsed Inner Membrane Complex (IMC) and unsegregated nuclei. **(a)** Confocal imaging of the Parental Tir1 and iKD TgPP1 strains labelled with TgEno2 (red) and TgIMC1 (green) in the presence and absence of auxin treatment. DAPI was used to stain the nucleus. Scale bar (1µm) is indicated in the lower right corner of each individual image. **(b)** Bar graph representing the percentage of Parental Tir1 and iKD TgPP1 vacuoles possessing a collapsed Inner Membrane Complex (IMC) by using anti-TgIMC1 antibodies for labelling the IMC in the absence and presence of auxin treatment for 48 hours. A Student’s *t*-test was carried out, ***p<0.001; mean ± s.d. (n=3). **(c)** Bar graph displaying the quantification of unsegregated nuclei in the Parental Tir1 and iKD TgPP1 strain in the absence and presence of auxin treatment for 48 hours. A Student’s *t*-test was carried out, ***p<0.001; mean ± s.d. (n=3). **(d)** Electron microscopy (EM) image demonstrating the structural morphology characteristics of the IMC and plasma membrane (PM) of the iKD TgPP1 mutant parasite in the absence of auxin treatment. In this case, both IMC and PM remain intact Magnified region is boxed. **(e)** EM image demonstrating the structural morphology of the IMC and PM of the iKD TgPP1 parasite after auxin treatment for 48 hours. In this case, PM membrane remains intact, but the IMC is absent. (I) stand for IMC. Scale bar (5 µm) is demonstrated in the lower right region of each EM. Magnified regions are boxed. Region missing the IMC are indicated by black arrows. A region with both the IMC and PM is indicated by a white arrow.

### The effect of TgPP1 depletion affects the division and segregation of the Golgi and Plastid

During the tachyzoite’s cell cycle, the sub-cellular organelles are replicated according to a well-determined timeline ^2^. Indeed, the centrosome divides first and is then followed by the Golgi and then Plastid ^21^. We investigated the effect of TgPP1 depletion on the division of the centrosome (using TgCentrin1 as a marker of the outer core centrosome) by IFA and observed that the shape of the centrosome is impacted by the depletion of TgPP1 (supplementary figure 3a, lowest panel). However, there was no change in the TgCentrin1 to nucleus ratio, suggesting that the centrosome division is unaffected in absence of TgPP1. We then investigated the effect of TgPP1 on the division of organelles. We, therefore, studied the effect of TgPP1 on the division of the plastid and the Golgi. By IFA, we observed that the plastid (TgCpn60) and Golgi (TgSORT) were missegregated after TgPP1 depletion (figure 3a). The depletion of TgPP1 after 48 hours of auxin treatment resulted in a significant percentage of parasite vacuoles with this phenotype (figure 3b). Collectively, these results demonstrate that TgPP1 has an important role in coordinating the cell-cycle steps after centrosome division.

**Figure 3:**
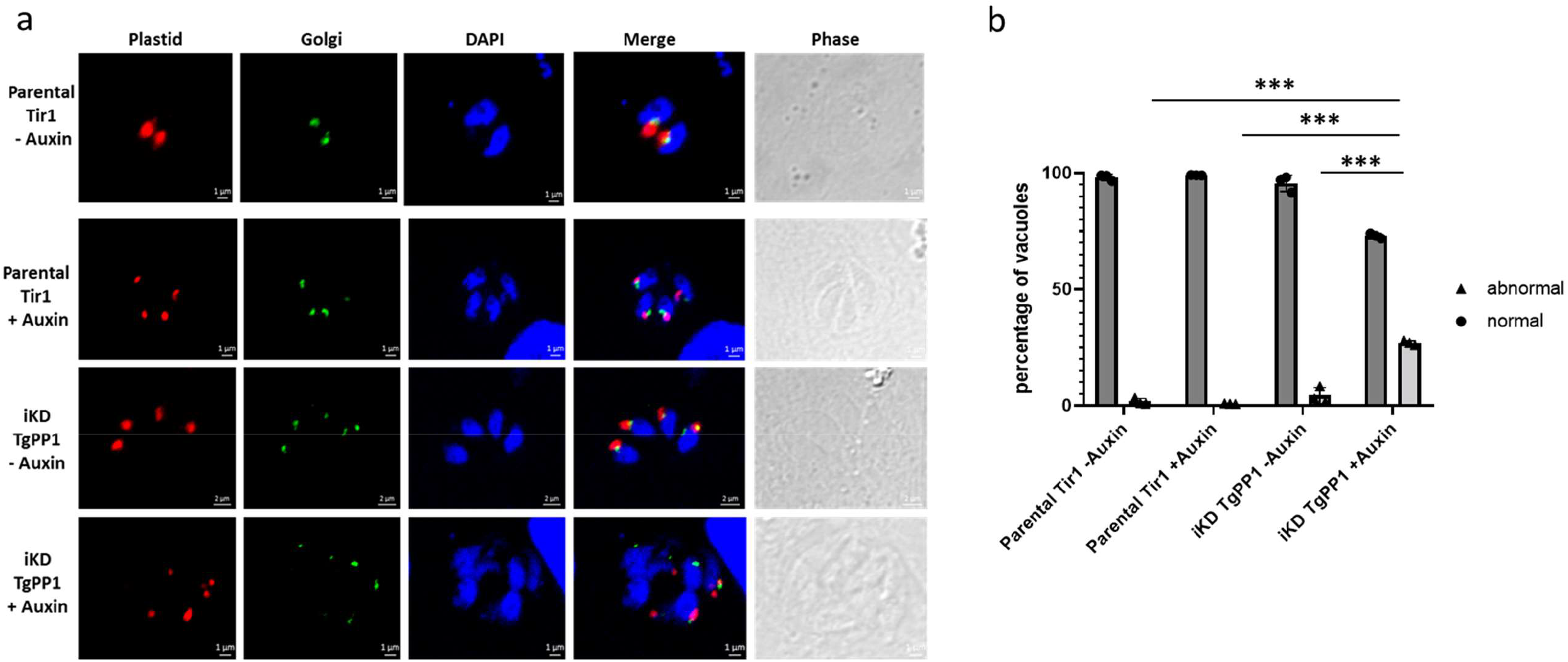
TgPP1 depletion induce plastid and Golgi missegregation. **(a)** Confocal imaging of the Parental Tir1 and iKD TgPP1 strains with labelled plastid (red) and Golgi (green) in the presence and absence of auxin treatment. DAPI was used to stain the nucleus. Scale bar (1µm) is indicated in the lower right corner of each individual image. **(b)** Graph bar demonstrating the percentage of Parental Tir1 and iKD TgPP1 vacuoles possessing normal and abnormal plastid and Golgi segregation in the absence and presence of auxin treatment for 48 hours. A Student’s *t*-test was carried out. ***p<0.001; mean ± s.d. (n=3). Dark grey bars represent normal plastid and Golgi segregation. Light grey bars represent abnormal plastid and Golgi segregation.

### TgPP1 depletion affects amylopectin steady-state levels

The control of amylopectin levels has been reported to be regulated by the phosphorylation ^13^ and dephosphorylation ^11,12^ of enzymes of the amylopectin metabolism in *T. gondii*. We hypothesized that TgPP1 could also play a role in starch storage. To measure the effect of TgPP1 depletion on the amylopectin steady-state levels, we used the Periodic Acid Shiff (PAS) method that labels polysaccharides. After 48 hours of TgPP1 depletion we clearly identified the appearance of structures positive for PAS staining accumulating in the parasites (figure 4a, lower panel). This labelling is almost inexistent in the parental strain (in presence or absence of auxin) but can be detected in the iKD TgPP1 mutant in absence of auxin (figure 4a). The presence of amylopectin granules was further confirmed through EM where they were absent in the parental strain in presence of auxin (figure 4b) but observed in the iKD TgPP1 parasite in the presence of auxin (figure 4c). The percentage of PAS positive vacuoles were quantified in the iKD TgPP1 mutant and parental strains in the absence and presence of auxin for 48 hours. There was a significantly increased percentage of PAS positive vacuoles (∼50%) in the iKD TgPP1 mutant in the presence of auxin when compared to the parental strain or the iKD TgPP1 mutant in absence of auxin (figure 4d). However, as observed in figure 4a, a significant number of vacuoles were PAS positive the iKD TgPP1 mutant in absence of auxin (around 15%). We independently verified the accumulation of amylopectin in the iKD TgPP1 mutant in absence and presence of auxin by a biochemical dosage of amylopectin produced by this strain and the parental strain. In the absence and presence of auxin, the iKD TgPP1 mutant parasite possesses a significant amount of amylopectin (figure 4e). The amount of amylopectin was significantly increased in presence of auxin in the iKD TgPP1 mutant (figure 4e), indicating that the depletion of TgPP1 induces the accumulation of amylopectin in the parasite. Therefore, TgPP1 plays pleiotropic roles during the intracellular growth of the tachyzoite including regulation of the amylopectin steady-state levels.

**Figure 4:**
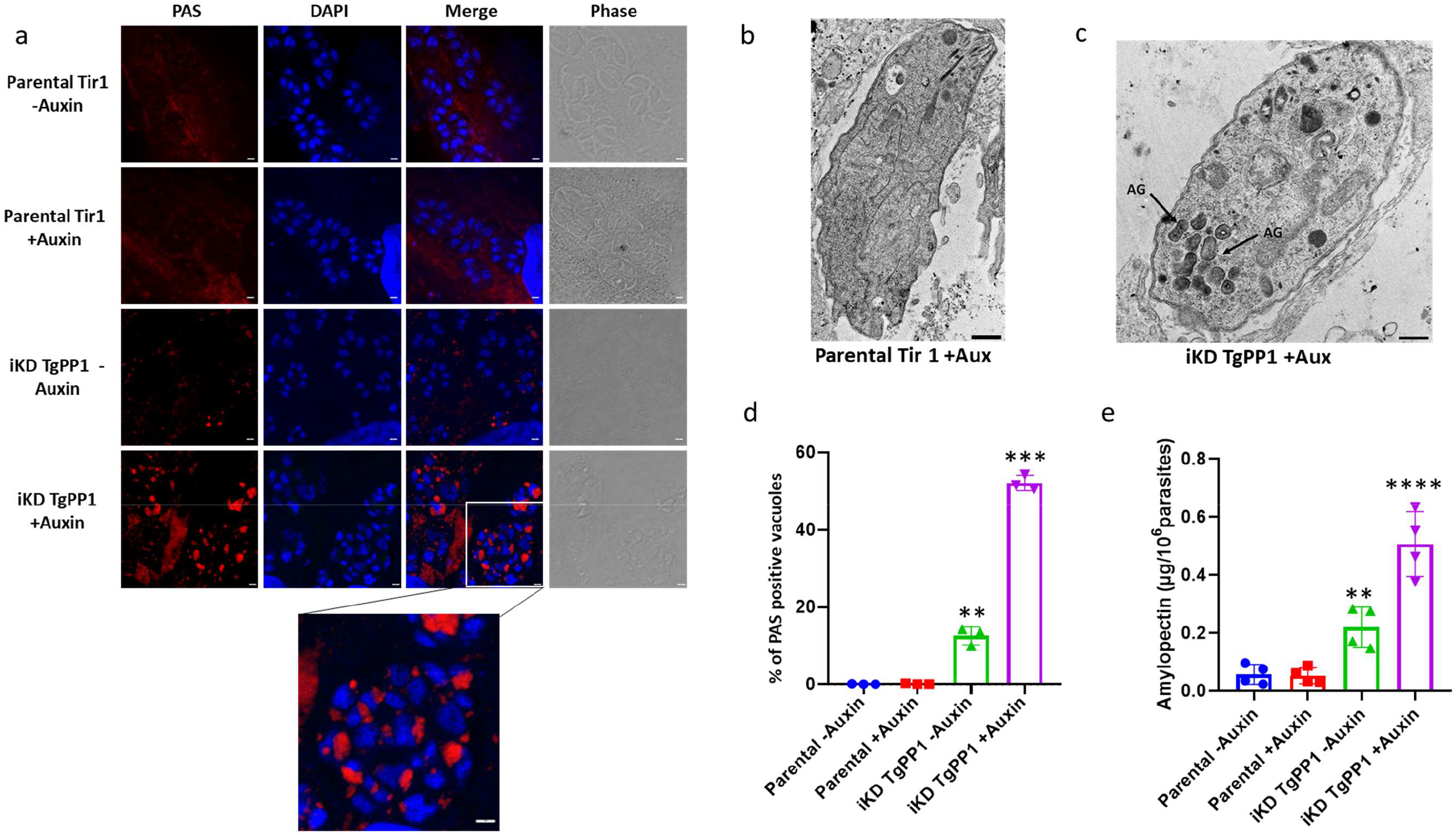
Absence of TgPP1 results in the accumulation of amylopectin granules. **(a)** Confocal imaging of the Parental Tir1 and iKD TgPP1 strain stained with PAS (red) in the absence and presence of auxin for 48 hours. DAPI was used to stain the nucleus. Scale bar (1µm) is located in the far-right corner of each image. **(b)** Electron microscopy scan of the Parental Tir1 and iKD TgPP1 parasite in the presence of auxin depicting the presence of amylopectin granules in the iKD mutant. Amylopectin granules (AG) are indicated by arrows. Scale bar (5µm) is demonstrated in the lower right region of each EM scan. **(c)** Bar graph representing the percentage of PAS positive vacuoles in the Parental Tir1 and iKD TgPP1 strains in the absence and presence of auxin for 48 hours. A Student’s *t*-test was carried out. **p<0.01, ***p<0.001; mean ± s.d. (n=3). **(d)** Quantification of the amount of amylopectin, as measured by biochemical assay, present within Parental Tir1 and iKD TgPP1 parasites in the absence and presence of auxin for 48 hours. A Student’s *t*-test was carried out. ***p<0.001, ****p<0.0001; mean ± s.d. (n=3).

### TgPP1 regulates the biology of the tachyzoite through dephosphorylation of a large set of proteins

Since TgPP1 was localized to the nucleus of the parasite, we investigated its potential role in gene regulation. For that we carried out RNA-sequencing analysis by examining changes in the transcriptome of the iKD TgPP1 strain after treatment with or without auxin for 24 hours from four biological replicates. Deseq2 was used to determine significant differential expression of genes based on an adjusted p-value cutoff of 0.05 and a minimum fold-change of 2 (supplementary figure 4a). To our surprise, only a few genes were considered differentially regulated following the depletion of TgPP1 and none of them pass the minimum fold-change of 2 cutoff (supplementary figure 4a, supplementary table 1). This suggested that TgPP1 had a minimal role in transcriptional regulation despite its nuclear localization.

We then investigated the role of TgPP1 in differential phosphorylation of proteins during the intracellular growth of the parasite. For that, the phosphoproteome of the iKD TgPP1 mutant parasite was investigated at two time points. A first time-point, after 24 hours of normal growth and then 2 hours of auxin treatment, was aimed at discovering the direct targets of TgPP1, a short time period after its complete depletion (as measured by Western-blots). A second time-point, after 24 hours of auxin treatment was aimed at measuring the global effect of TgPP1 on the phosphoproteome. The presence of significantly hyper or hypo-phosphorylated peptides was identified in a minimum of three biological replicates.

After 2 hours of auxin treatment, a total of 7822 phosphorylation sites were identified (supplementary table 2), among the 4 biological replicates that were analyzed. Among these, 104 phosphopeptides exhibited significant differential phosphorylation with an FDR less than 0.01 (figure 5a and supplementary table 2). Among the 104 phosphopeptides (representing 93 proteins), 37 proteins were considered significantly hyper-phosphorylated (40 phospho-peptides associated) and 56 proteins (64 phospho-peptides associated) were considered significantly hypo-phosphorylated (figure 5a). We focused on the hyper-phosphorylated proteins that are likely the effect of the phosphatase depletion. We noticed that IMC1 was identified with the highest hyper-phosphorylation ratio (supplementary table 2). Moreover, the Apical Cap protein 2 (AC2, TGME49_250820), the CPH1-interacting protein 2 (CIP2, TGME49_257300) and TGME49_285850 are predicted to localize to the apical compartment or IMC ^22^ and were found to be hyperphosphorylated after 2h of auxin treatment. Concordant with the TgPP1 nuclear localization, a high number of hypo-phosphorylated proteins (19/37) are predicted to localize to the nucleus. Among them, the DNA replication licensing factor MCM7 is likely involved in DNA replication. We also identified hyper-phosphorylation of two fitness-conferring kinases: CDPK6^23^ and TKL1 ^24^. Moreover, we identified the hyper-phosphorylation of a trehalose phosphatase (TGGT1_297720), an uncharacterized protein containing a CBM20 domain, known to bind to starch. In the list of hypo-phosphorylated proteins, we also found 5 other IMC proteins such as IMC17, ISC1, PMCAA1, and uncharacterized IMC proteins (TGGT1_217510 and TGGT1_306190). Moreover, 4 putative kinases (CDPK2a, the cell cycle associated GSK (TGME49_265330), TGME49_225960 and TGME49_320000) and one phosphatase (TGME49_269460) were also found hypo-phosphorylated.

**Figure 5:**
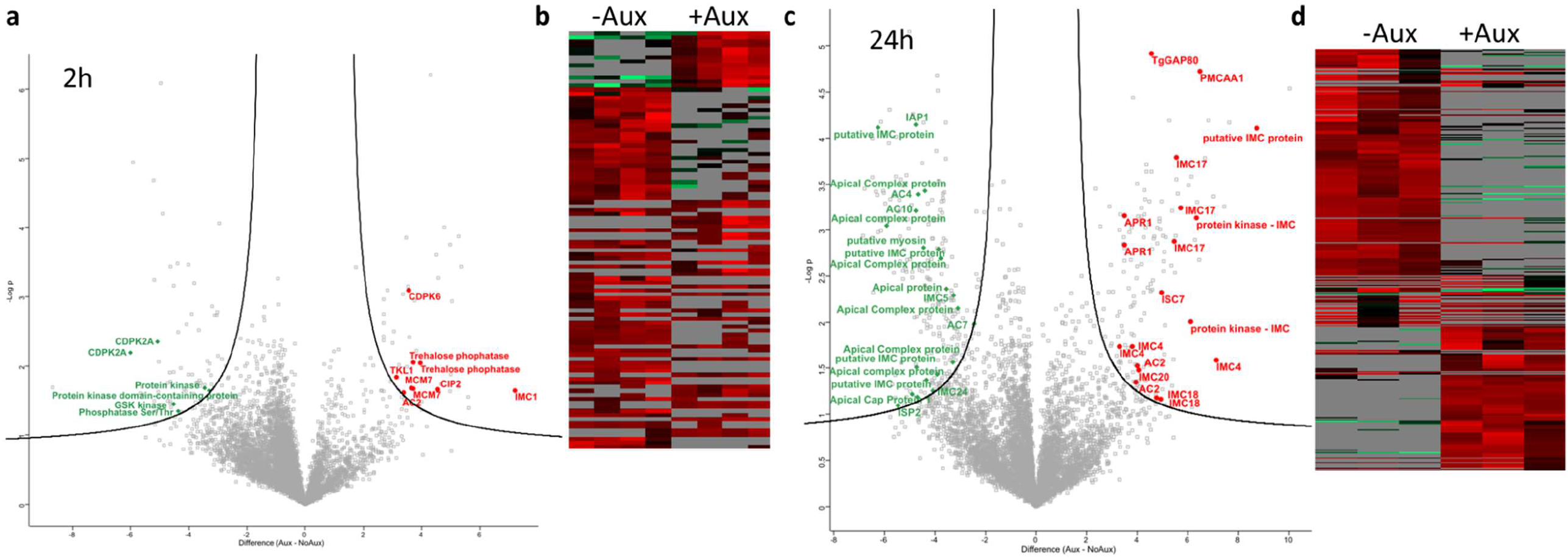
TgPP1 depletion results in differentially phosphorylated IMC proteins. **(a)** Volcano plot of the total phosphosites in the iKD TgPP1 mutant parasite after the treatment of auxin for 2 hours resulting from phosphoproteomics analysis (n=7822). Selected proteins presenting hyperphosphorylated peptides in absence of TgPP1 are highlighted in red. Selected proteins presenting hypophosphorylated peptides are highlitted in green. See the full list in supplementary Table 2. **(b)** Heat map of phosphosites which are differentially phosphorylated as a result of 2 hours of auxin treatment in the iKD TgPP1 mutant. **(c)** Volcano plot of the total phosphosites in the iKD TgPP1 mutant parasite after the treatment of auxin for 24 hours as demonstrated following phosphoproteomics analysis (n=7509). Selected proteins presenting hyperphosphorylated peptides in absence of TgPP1 are highlighted in red. Selected proteins presenting hypophosphorylated peptides are highlitted in green. See the full list in supplementary Table 3. **(d)** Heat map displaying phosphosites which are differentially phosphorylated following the treatment of auxin for 24 hours.

We then wanted to investigate further by studying the effect of TgPP1 depletion on the differential phosphorylation of phospho-peptides after a longer period of auxin treatment. After 24 hours of auxin treatment, a total of 7509 phospho-sites were identified. Out of these phospho-sites, 376 were considered as significantly differentially phosphorylated and corresponded to a total of 284 proteins (figure 5b and supplementary table 3). The total number of hyper-phosphorylated peptides was 152 whereas the total number of hypo-phosphorylated peptides was 224, associated with 114 and 178 proteins, respectively. A striking number of IMC proteins (10/114) were discovered to be significantly hyper-phosphorylated such as IMC4, IMC18, IMC20, AC2 and ISC7. On the other hand, 17 proteins associated with the IMC or the apical compartment were identified as hypo-phosphorylated, suggesting a global perturbation of the phosphorylation status of this compartment. As an illustration, a single IMC protein, IMC17, was found both hyper-phosphorylated at particular phospho-sites and hypo-phosphorylated at other phospho-sites. Of note, the Trehalose phosphatase (TGME49_ 297720) was also found hyper-phosphorylated after 24 hours of treatment.

Of note, we performed a global proteome analysis of the same samples that were treated for 24h with or without auxin (Supplementary Table 4 and Figure S4b) and did not identify significant differences in protein content in these samples, suggesting that TgPP1 mainly acts through modulating the phosphorylation status of proteins and does not affect global proteome content.

### TgPP1 dephosphorylation of the TGME49_297720 protein may influence starch metabolism

TgPP1 depletion influences starch metabolism (figure 4). We identified the two differentially phosphorylated sites on the TGME49_297720 protein. This protein encodes for a starch binding domain (CBM20), a trehalose-6-phosphate synthase (TPS), and a trehalose-6-phosphate phosphatase (TPP) domain (figure 6a). Both identified hyper-phosphorylated sites (Serine 1054 or Serine 1073) are placed outside of the putative enzymatic domains (figure 6a). We explored whether these phospho-sites were significantly linked to the starch accumulation phenotype observed previously. For that, we produced four transgenic parasite mutants bearing single point mutations corresponding to Serine 1054 or Serine 1073 changed to an Alanine or Aspartate amino acid. A parental strain (RH ΔKu80) with the Trehalose phosphatase gene bearing a C-terminal Myc tag was used to generate the mutants using the CRISPR/Cas9 system and direct FACS sorting into 96 well plates. To identify whether the WT or mutated fragment had been inserted into the correct locus of the genome, PCR fragments representing the targeted locus were amplified and sent for sequencing (supplementary figure 5a). First, we verified that the mutation had no effect on the enzyme localization (Figure 6b). We then proceed to verify the effect of the induced mutation on accumulation of starch in these mutant parasites. For that, we quantified the number of PAS positive vacuoles in normal culture conditions (pH 7) or in conditions known to induce bradyzoite differentiation (pH 8.2, the bradyzoite form accumulates amylopectin. In normal culture conditions, there was no significant difference between the WT and the mutant parasites, when measuring the percentage of PAS positive vacuoles. However, in bradyzoite inducing conditions (pH 8.2), the percentage of PAS positive vacuoles significantly decreased in the 1073A mutant whereas it significantly increased in the 1073D mutant compared to their wild-type counterpart. These results suggest that the phosphorylation status of the Trehalose phosphatase is linked to the ability of the parasite to accumulate starch, mimicking the phenotype observed after TgPP1 depletion.

**Figure 6:**
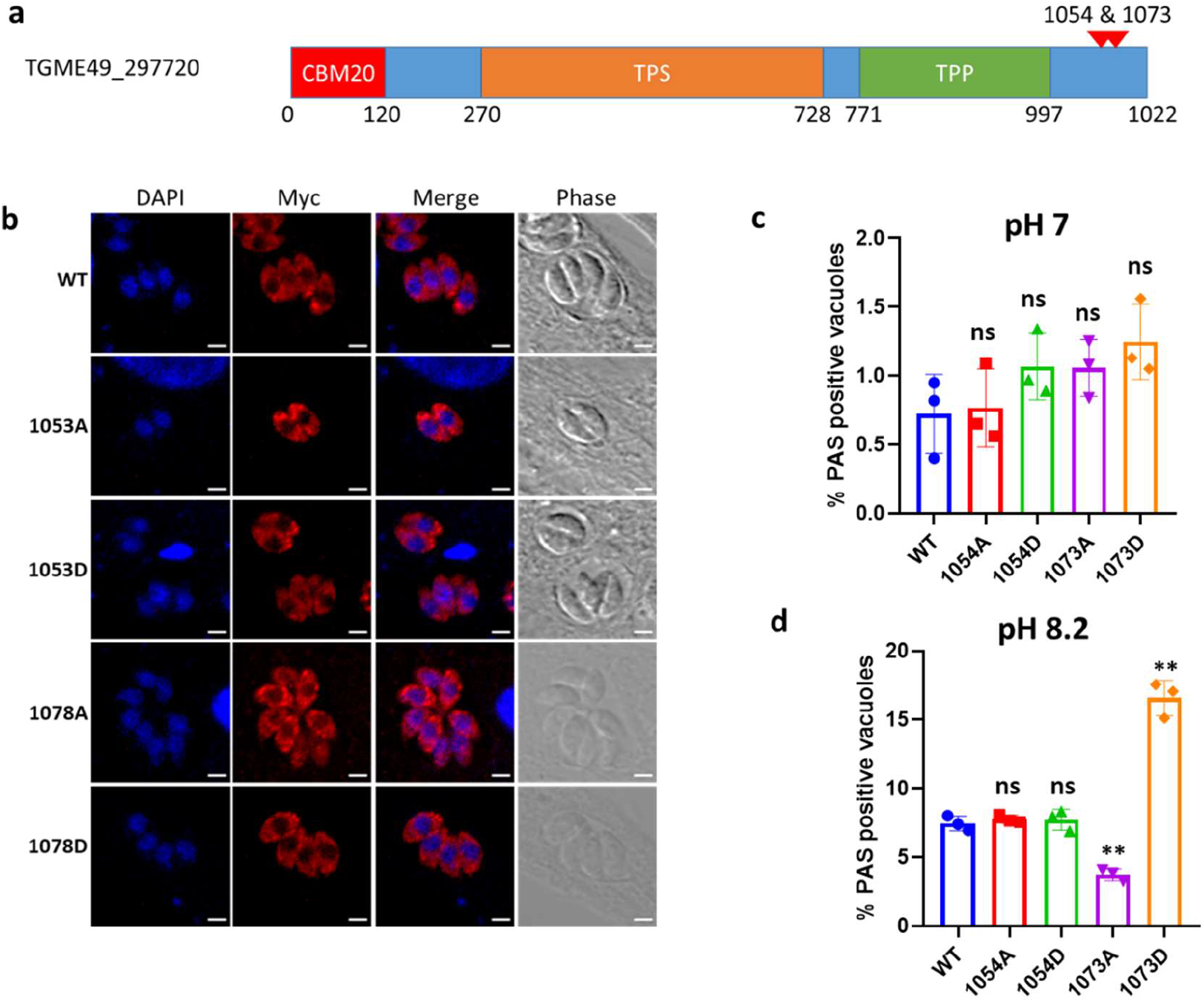
The phosphorylation status of the TGGT1_297720 protein affects amylopectin steady-state levels. **(a)** Schematic representation of the trehalose synthase-phosphatase protein (TGGT1_297720) that encompasses consisting of CBM20, Trehalose synthase (TPS), and the Trehalose phosphatase (TPP) domains. Differentially phosphorylated site after TgPP1 depletion at Serine 1054 and Serine 1073 are indicated by red arrow heads. **(b)** Confocal imaging of the mutated TGGT1_297720 proteins. The myc tagged TGGT1_297720 protein the labelled with anti-myc antibodies (red). DAPI was used to stain the nucleus. Scale bar (2 µm) is indicated in the lower right corner of each individual image. The identity of the mutation is indicated on the left side. **(c)** Bar graph representing the percentage of PAS positive vacuoles at a pH of 7.0 in the WT and mutant 1054A, 1054D, 1073A, and 1073D parasites. A Student’s *t*-test was carried out. ns>0.05; mean ± s.d. (n=3). **(d)** Bar graph representing the percentage of PAS positive vacuoles at a pH of 8.2 in the WT and mutant 1054A, 1054D, 1073A, and 1073D parasites. A Student’s *t*-test was carried out. ns> 0.05, **p <0.01; mean ± s.d. (n=3).

## DISCUSSION

Phosphorylation is a widespread post-transcriptional modification in apicomplexan parasites ^5^. In *T. gondii*, it is involved in the regulation of the tachyzoite cell cycle ^21^ and of the starch metabolism^13^. Here, we explored the pleiotropic phenotypes caused by the depletion of the TgPP1 protein. TgPP1 is involved in a wide range of molecular mechanisms including nuclear and organelle segregation. TgPP1 is a crucial phosphatase that regulates both the intracellular cell cycle and the extracellular mobility ^14^. Genetic manipulation of this locus has proven challenging and the resulting independent mutants have shown a decreased fitness in conditions where TgPP1 was still expressed suggesting that the activity of this protein is highly important for the parasite. The mutant produced in this study, although noticeably affected in absence of auxin, showed stronger phenotypes under auxin treatment as shown by the plaque assay performed.

Among the phenotype observed, it was affected in IMC formation, a phenotype that was confirmed in IFA and EM. Differential phosphorylation in absence of TgPP1 of multiple IMC targeted proteins such as IMC1, IMC4, IMC5, IMC17, IMC18, IMC20, IMC24, ISP1, and ISC1 is a hallmark of the phospho-proteomics experiments we performed. A short time after TgPP1 depletion, TgIMC1 is found to be hyper-phosphorylated at the threonine 62, with the greatest amplitude when comparing the auxin treated samples to the non-treated ones. IMC proteins are targeted sequentially to the IMC and form a rigid meshwork onto the flattened alveolar sacs ^25^. Post-translational modifications can influence the function and assembly of the IMC proteins. For example, TgIMC1 undergoes proteolytic cleavage and the cleaved form is associated with filament rigidity ^26^. Numerous phosphorylation sites have been detected on IMC proteins although their biological significance remained to be investigated. Of note, assembly and rigidity of intermediate filament proteins (e.g., nuclear lamina network) in other organisms have been shown to be regulated through phosphorylation ^27^. Phosphorylation on serine and threonine residues would generally promote disassembly and dephosphorylation increasing stability of the intermediate filament ^28^. After TgPP1 depletion, we have found a global collapse of the IMC and hyper-phosphorylation of multiple IMC proteins that suggest that the IMC network assembly and stability may be controlled by similar phosphorylation mechanisms as for the assembly of other intermediate filaments. The specific role in the assembly of the IMC of the TgIMC1 Thr62 phosphorylation would be of interest to investigate in the future, although individual phosphorylation sites on TgIMC12 were not shown to impact the IMC assembly ^29^. Rather than the limited phosphorylation status of individual IMC proteins, TgPP1 depletion seems to induce a global imbalance in the phosphorylation status of IMC proteins that may be the cause of the visual phenotype observed.

Phosphorylation of TgGAP45 has been shown to control its assembly to the MyoA-glidosome^30^, a structure essential for the extracellular movement of the parasite. Interestingly, TgGAP80, a protein belonging to the TgGAP45 protein family, was found to be hyperphosphorylated after TgPP1 depletion. Moreover, IAP1, a protein associated with TgGAP80 is also differentially phosphorylated in presence of auxin. Both proteins are part of the MyoC-glidosome, a complex associated with IMC proteins that localizes at the basal pole of the parasite ^31^. Phosphorylation of the GAP45 family of proteins might be therefore a general mechanism that ensures their association with the component of the glidosome. These data indicate that the phosphorylation status of IMC proteins might be an important determinant of their ability to assemble and rigidify the IMC network. The structural morphology of the iKD TgPP1 mutant is impacted as was indicated through EM. In the presence of auxin, the usual bow-shaped tachyzoite is absent and instead an abnormally shaped tachyzoite takes its place. Despite that the plasma membrane remains intact, it appears to exhibit unusual curvatures and invaginations, this might possibly be due to the collapsed IMC structure which is absent underneath it. In normal conditions, the tachyzoite structure is maintained through IMC proteins resulting in tachyzoite pellicle strength.

The presence of nuclear and organelle segregation phenotypes after depletion of TgPP1 suggests that this phosphatase may have targets that are organizing cell division in the parasite. Although the centrosome division remains unaffected in absence of TgPP1, an obvious target would be the centrosome complex. However, none of the known centrosomal proteins were identified in the phospho-proteomics analysis. Although, the analysis performed here is far from being exhaustive, a protein of unknown function targeted by TgPP1 may have centrosomal localization. Indeed, the centrosome composition remains only partially known in *T. gondii* ^32^. Overall, the large changes in the phosphorylation status of kinases may also affect the centrosome function, although we were not able to pinpoint one known centrosomal regulator.

RNA-seq revealed that TgPP1 depletion did not change the transcriptome of the parasite after 24 hours of auxin treatment. Despite TgPP1 nuclear localization and the presence of differential phosphorylation of ApiAP2 transcription factors in the phospho-proteome at 24h auxin treatment, the phenotype observed are not the consequence of differential transcript expression or proteome expression. ApiAP2 post-transcriptional modifications have been suggested to modify their activity ^2^ but the data presented here does not support this hypothesis.

The formation of amylopectin granules in the tachyzoite following the depletion of TgPP1 was a striking finding. This suggests that TgPP1 has a role in the regulation of amylopectin steady-state levels during tachyzoite proliferation. Phosphorylation has been shown to regulate amylopectin metabolism in *T. gondii*. TgCDPK2 phosphorylate a wide range of enzymes that are involved in this process ^13^ among which a glycogen phosphorylase, an alpha-glucan water dikinase, and a pyruvate phosphate dikinase that are known to bind to amylopectin through CBM20 domains ^33^. Mutating the phosphorylation sites of glycogen phosphorylase was later shown to phenocopy the TgCDPK2 knock-out strain ^34^ suggesting a crucial role for phosphorylation in regulating the steady-state levels of amylopectin in this parasite. A phosphatase was also recently implicated in this pathway: TgPP2A contributes to the regulation of amylopectin metabolism via dephosphorylation of TgCDPK2 at a particular site (S679) ^11^ suggesting a pivotal role for TgCDPK2 in amylopectin accumulation. We found that a protein annotated as a Trehalose phosphatase was differentially phosphorylated in absence of TgPP1. Mutagenesis of these phosphosites recapitulates partially the TgPP1 depletion phenotype with amylopectin accumulation being detected after induction of bradyzoite differentiation. Interestingly, the TGME49_297720 protein was not listed as the potential target of TgCDPK2 or TgPP2A suggesting that it acts through an independent pathway that also controls the metabolism of amylopectin. This protein encompasses a starch-binding domain (CBM20) that further supports the hypothesis of a function linked to starch metabolism regulation. This protein is annotated as a Trehalose phosphatase. However, its sequence encodes a trehalose-6-phosphate synthase (TPS) and a trehalose-6-phosphate phosphatase (TPP) domain. Dual activity enzymes are found in some bacteria and fungi ^35^. The TGME49_297720 protein may therefore be misannotated as trehalose phosphatase and should be annotated as a bifunctional TPS-TPP protein. The presence of this dual-activity enzyme indicates that *T. gondii* may have the ability to produce trehalose from UDP-Glucose and Glucose-6-Phosphate as it is the case for numerous eukaryotes. However, these activities have to be confirmed experimentally. Trehalose is a non-reducing disaccharide consisting of two glucose subunits with an α,α′-1,1′-glycosidic bond. This carbohydrate occurs in a wide range of species and is synthesized by bacteria, fungi, plants, and various invertebrates. Trehalose has roles in development, stress tolerance, energy storage, and the regulation of carbon metabolism in plants and fungi ^36^. While trehalose has not been identified so far in the parasite ^37^, the data presented here suggest that it may have a role in regulating the starch metabolism in the parasite.

Overall, we showed that TgPP1 acts through the phosphorylation of a wide range of targets to regulate a large set of molecular mechanisms that are essential for the completion of the tachyzoite cell cycle. Our study put an emphasis on the role of phosphorylation in the IMC protein network assembly and stability. It also confirms the importance of phosphorylation in regulating the amylopectin steady-state levels in the parasite and suggests the involvement of a new regulatory pathway involving a new enzyme in this pathway.

## Data Availability Statement

The mass spectrometry proteomics data have been deposited to the ProteomeXchange Consortium via the PRIDE partner repository with the dataset identifier PXD043539. The RNA-seq data was deposited to the SRA database under the identifier PRJNA1002988.

