## Supplemental figures 1-4 for "The PP1 phosphatase exhibits pleiotropic roles controlling both the tachyzoite cell cycle and amylopectin-steady state levels in *Toxoplasma gondii*"

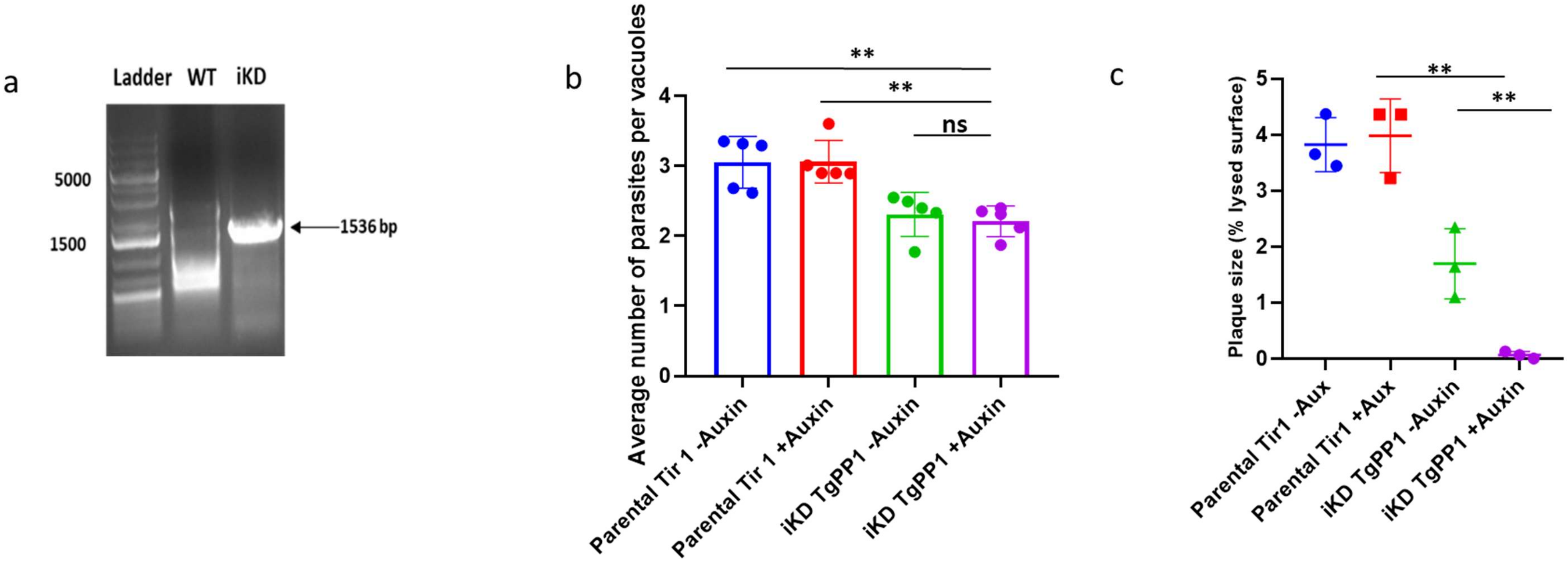


**Supplementary Figure 1: iKD TgPP1 mutant construction (a)** PCR verifying integration of the HXGPRT-2TA-AID-Ty cassette at the correct genome locus of the iKD TgPP1 mutant. A band corresponding to 1536 using iKD TgPP1 genomic DNA confirms cassette integration compared to using WT genomic DNA. **(b)** Growth assay of the Parental Tir1 and iKD TgPP1 mutant strains in the absence and presence of auxin treatment for 24 hours. A Student's *t*-test was performed, ns>0.05, **p<0.01; mean ± s.d. (n=5). **(c)** Bar graph indicating plaque size produced by the Parental Tir1 and iKD TgPP1 strain in the presence and absence of auxin. Plaque size was determined by measuring the percentage of lysed surface of the plaque assay. Three independent experiments were carried out. A Student's *t*-test was performed, **p<0.01; mean ± s.d. (n=3).

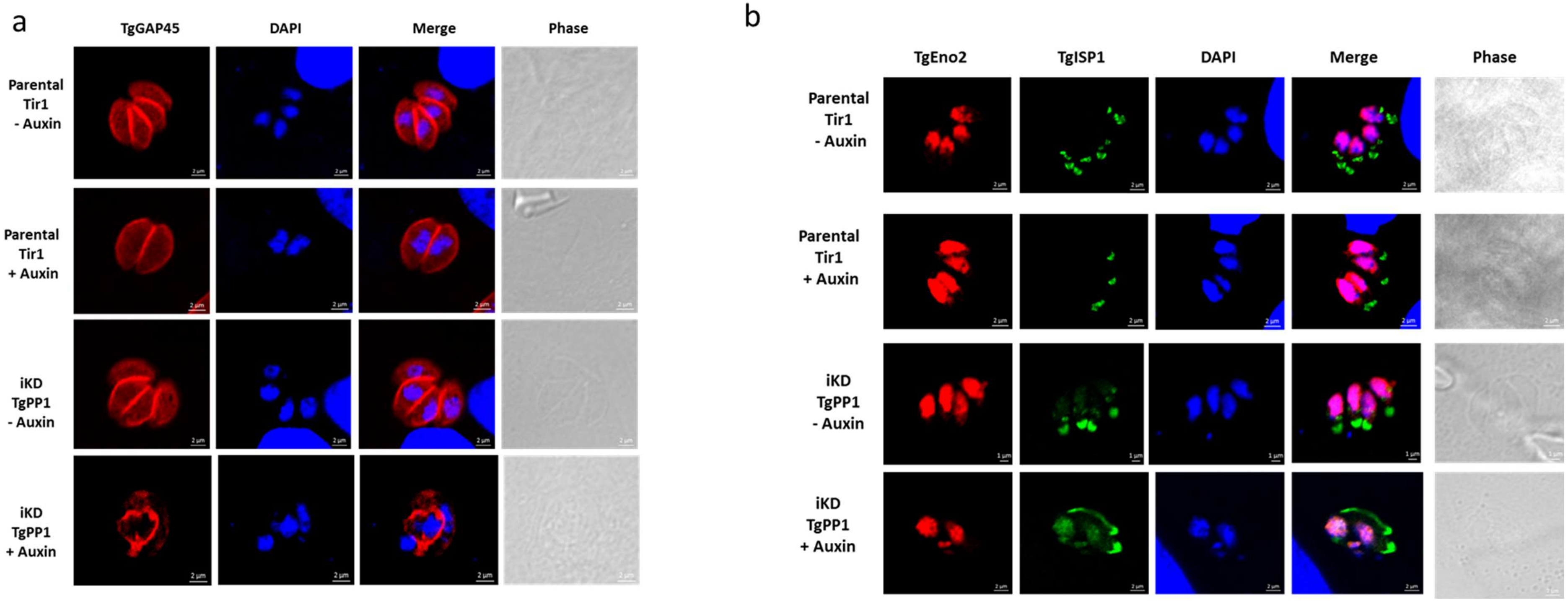


**Supplementary Figure 2: iKD TgPP1 demonstrates a collapsed IMC verified through TgGAP45 and TgISP1 labelling (a)** Confocal imaging of the Parental Tir1 and iKD TgPP1 strains labelled with anti-TgGAP45 (red) in the presence and absence of auxin treatment. DAPI was used to stain the nucleus. Scale bar (1µm) is indicated in the lower right corner of each individual image. **(b)** Confocal imaging of the Parental Tir1 and iKD TgPP1 strains labelled with anti-TgEno2 (red) and anti-TgISP1 (green) in the presence and absence of auxin treatment. DAPI was used to stain the nucleus. Scale bar (1µm) is indicated in the lower right corner of each individual image.

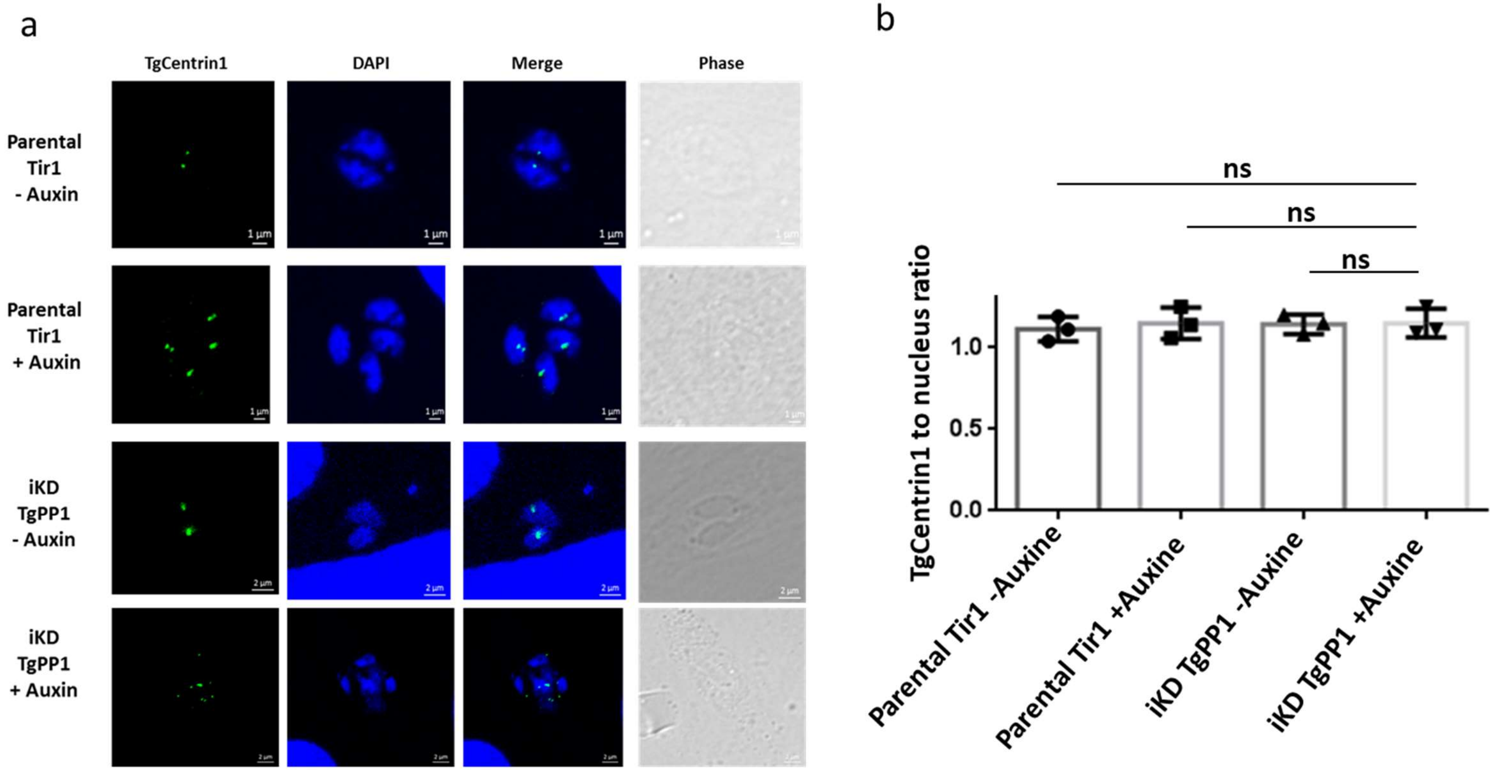


**Supplementary Figure 3: Conditional depletion of TgPP1 has a qualitative impact on the outer core centrosome (a)** Confocal imaging of Parental Tir1 and iKD TgPP1 parasites in the absence and presence of auxin treatment for 48 hours labelled with anti-TgCentrin1 antibodies (green). DAPI was used to stain the nucleus. Scale bar (1µm) is indicated in the lower right corner of each image. **(b)** Bar graph demonstrating TgCentrin1: nucleus ratio of Parental Tir1 and iKD TgPP1 in the absence and presence of 48-hour auxin treatment. A Student's *t*-test was performed, ns $p>0.05$; mean ± s.d. (n=3).

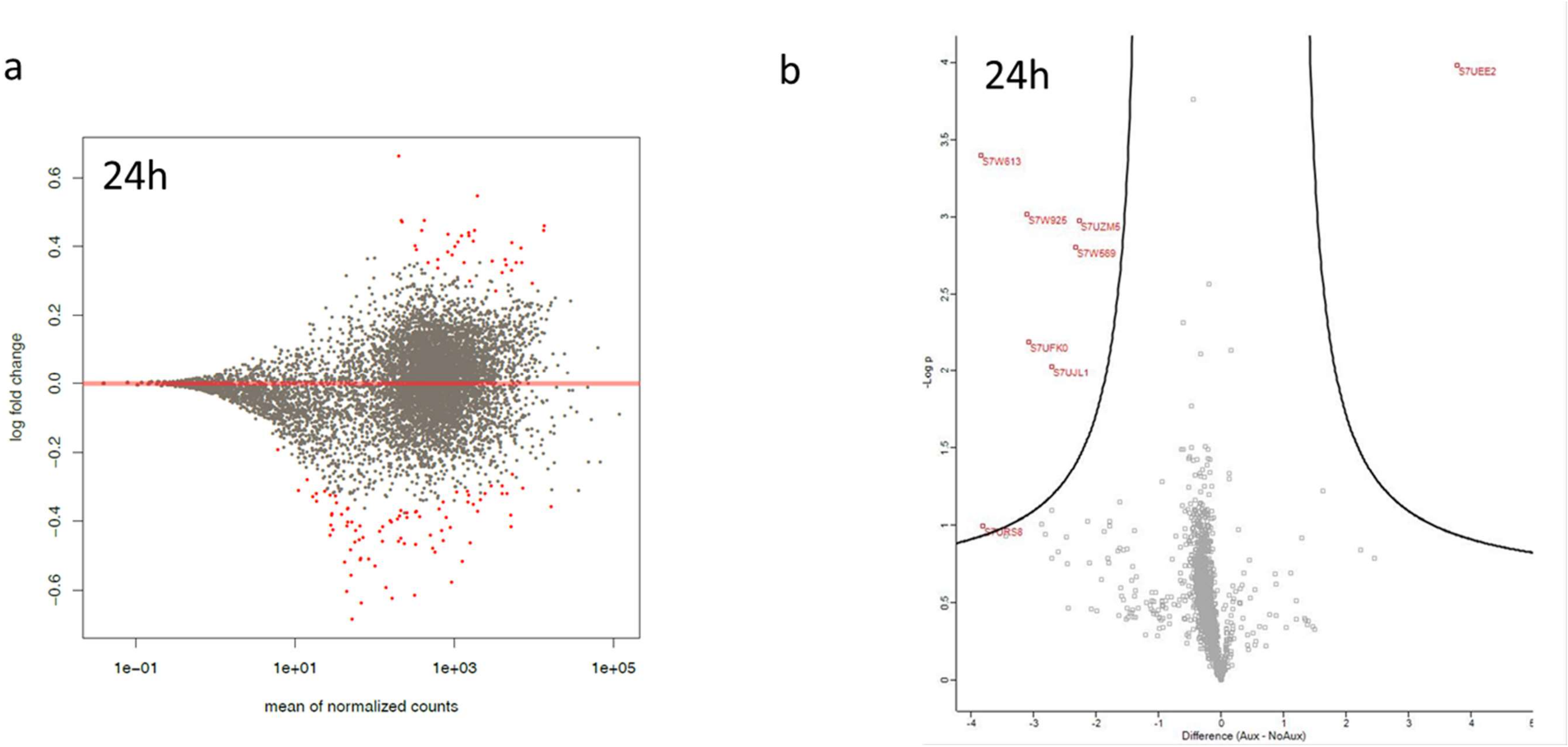


**Supplementary Figure 4: RNA-sequencing and proteome analysis at 24 hour of auxin treatment displays only a few differentially regulated genes (a)** Volcano plot demonstrating the differentially expressed genes analyzed from RNA-sequencing of the iKD TgPP1 mutant parasite treated with auxin for 24 hours. Differential expression is based on the analysis of three biological replicates. Statistically significant differentially expressed genes are indicated in red. However, these do not pass the +/- 1 $\log_2$ expression ratio criteria. **(b)** MA (Bland-Altman) plot demonstrating the differential proteome content in iKD TgPP1 mutant parasites after 24 hours of auxin treatment compared to iKD parasite grown in the absence of auxin (control). Statistically significant differentially expressed proteins are indicated in red.
